# Physical limit to concentration sensing in a changing environment

**DOI:** 10.1101/733840

**Authors:** Thierry Mora, Ilya Nemenman

## Abstract

Cells adapt to changing environments by sensing ligand concentrations using specific receptors. The accuracy of sensing is ultimately limited by the finite number of ligand molecules bound by receptors. Previously derived physical limits to sensing accuracy have assumed that the concentration was constant and ignored its temporal fluctuations. We formulate the problem of concentration sensing in a strongly fluctuating environment as a non-linear field-theoretic problem, for which we find an excellent approximate Gaussian solution. We derive a new physical bound on the relative error in concentration *c* which scales as *δc*/*c* ~ (*Dacτ*)^−1/4^ with ligand diffusivity *D*, receptor cross-section *a*, and characteristic fluctuation time scale *τ*, in stark contrast with the usual Berg and Purcell bound *δc*/*c* ~ (*DacT*)^−1/2^ for a perfect receptor sensing concentration during time *T*. We show how the bound can be achieved by a simple biochemical network downstream the receptor that adapts the kinetics of signaling as a function of the square root of the sensed concentration.

---

Cells must respond to extracellular signals to guide their actions in the world. The signals typically come in the form of changing concentrations of various molecular ligands, which are conveyed to the cell through ligand binding to cell surface receptors. A lot of ink has been expended on deriving the fundamental limits to the precision with which a cell can measure the concentrations from the activity of its receptors, constrained by the stochasticity of ligand binding and unbinding [1–4]. In particular, it has become clear that the temporal sequence of binding-unbinding events carries more information about the underlying ligand concentration than just the mean receptor occupancy, typically used in deterministic chemical kinetics models of this problem [5]. In particular, such precise temporal information allows cells to estimate the concentration of a cognate ligand even in a sea of weak spurious ligands [6–8], as well as to estimate concentrations of multiple ligands from fewer receptor types [9, 10], and molecular network motifs able to perform such complex estimation exist in the real world, even potentially taking advantage of cross-talk between receptor-ligand pairs [11].

Importantly, concentrations of ligands are worth measuring only when they are *a priori* unknown; or, in other words, if they change with time, allowing for instance cells to adapt their behaviour accordingly and maximize their long-term growth [12]. However, all of the preceding analyses have focused on the regime with a clear time scale separation, where the concentration is constant or constantly changing [13] during the period over which it is estimated. In this article, we will fill in this gap by calculating the accuracy with which a temporally varying ligand concentration may be estimated from a sequence of binding and unbinding events. This requires making assumptions about the time scale over which significant changes of the concentration are possible. In our formulation, the optimal sensor performs a Bayesian computation, formalized mathematically as a stochastic field theory. Crucially, we show how simple biochemical circuits allow one to perform the relevant complex computations.

## Field theory of concentration sensing

We associate to the ligand concentration *c*(*t*) a field *φ*(*t*) through *c*(*t*) = *c*_0_*e*^−*φ*(*t*)^, where *c*_0_ is an irrelevant reference concentration. Ligand concentration controls the ligand-receptor binding rate *r*(*t*) = 4*Dac*(*t*) = 4*Dac*_0_*e*^−φ(*t*)^ ≡ *r*_0_*e*^−φ(*t*)^, where 4*Da* is the diffusion-limited binding rate per molecule of the ligand to its target receptor, modeled as a circle of diameter *a* on the cell’s surface, and *D* is the ligand diffusivity. This binding rate can be readily generalized to *N* receptors by using instead *r*(*t*) = 4*N Dac*(*t*). All our results will then hold with this additional *N* factor. We assume that the concentration follows a geometric random walk, with characteristic time scale *τ*: *dφ* = *τ*^−1/2^*dW*, with *W* a Wiener process. This choice is justified by the fact that in many biological contexts, such as bacterial chemotaxis, concentrations may vary over many orders of magnitude.

The probability of the concentration temporal evolution over the time interval (0, *T*) is given by

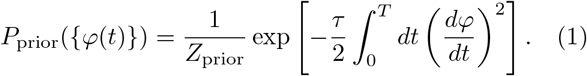

The receptor sees binding events at times *t*_1_, *t*_2_, …, *t_n_*, each occuring with rate 4*Dac*(*t_i_*) = *r*_0_*e*^−*φ*(*t_i_*)^. To simplify, let us assume that unbinding is instantaneous (generalization to finite binding times is discussed later). The posterior distribution of the concentration profile then follows Bayes’ rule:

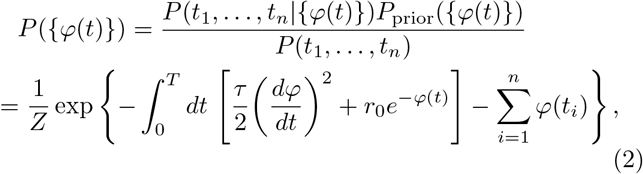

where *Z* is a normalization constant independent of *φ*. The term *r*_0_*e*^−*φ*^*dt* in the integral corresponds the probability of not binding a ligand between *t* and *t* + *dt* (except at times *t_i_*). The binding events at *t* = *t_i_* are generated by the *true* temporal trace of ligand concentration, *c*\*(*t*) = *c*_0_*e*^−*φ*\*(*t*)^. In the following the true trace *φ*\*(*t*) will be distinguished from the field *φ*, which refers to our observation-based belief.

The one-dimensional field-theoretic problem (2) is a particular case of Bayesian filtering [14]. When collecting information from binding events, cells do not have access to the future and cannot use the full span [0, *T*] of observations to infer the concentration at time *t*. Instead, they must infer it solely based on past observation in the interval [0,*t*], which distinguishes our problem from the mathematically similar inference of a continuous probability density [15–19]. This inference can be performed recursively by the rules of Bayesian sequential forecasting, similar to the transfer matrix technique, and also known as the forward algorithm [14]. To do this recursion, we first define:

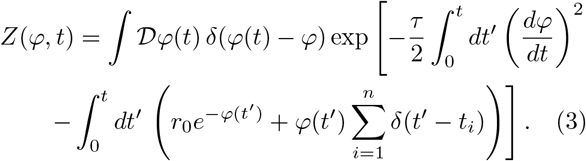

Considering past observations during the interval [0,*t*], the posterior distribution of *φ* at time t reads:

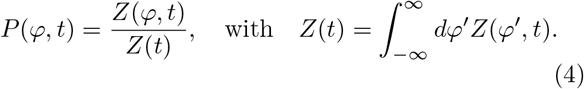

When considering periods during which no binding event was observed, we can write a recursion for *Z*(*φ*, *t*) between *t* and *t* + *dt*. Taking the *δt* → 0 limit yields, for *t* ≠ *t_i_* (App. A [20]):

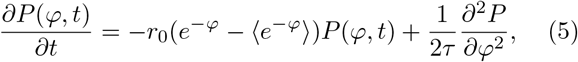

where 〈·〉 denotes an average over *P*(*φ*). When a binding even does occur at time *t_i_*, the posterior distribution is updated using Bayes’ rule:

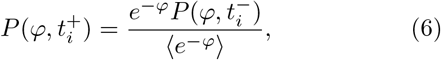

where 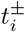 refer to the values right before and after the observation. The partition function *Z*(*t*) can be similarly calculated (App. A [20]) and could in principle be used to infer the correct timescale *τ* by maximizing *P*(*τ*|{*t*_1_, …, *t_N_*}) ∝ *Z* (App. C [20]).

## Gaussian solution

Because of the *P*(*φ*) dependence in 〈*e*^−*φ*^〉, the equations for the evolution of the posterior probability (5)-(6) are nonlinear. However, assuming a Gaussian Ansatz 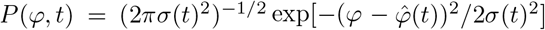, which is accurate in the limit of long measurement times (see below), gives a closed-form solution (App. B [20]), with:

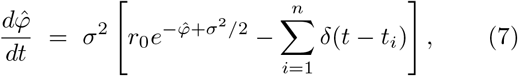

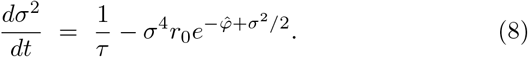

The maximum a posterior estimator for the concentration is then simply given by 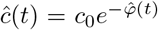, while *σ*(*t*)^2^ defines the Bayesian uncertainty on the estimator.

To check the validity of the Gaussian solution, we simulated (5)-(6) numerically, starting from a uniform distribution (*P*(*φ*, 0) = 1/2 for *φ* ∈ [−1,1] and 0 otherwise), with *r*_0_*τ* = 50 and a true *φ*\*(*t*) starting at *φ*\*(0) = 0. The numerical solution quickly approaches the Gaussian solution given by (7)-(8) starting with 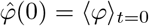 and *σ*(0)^2^ = Var(*φ*)_*t*=0_. The Kullback-Leibler divergence between the numerical and analytical solutions falls rapidly (Fig. 1A) and the numerical solution approaches the predicted Gaussian very closely (Fig. 1A, inset). Thus, the Gaussian solution provides an excellent approximation.

**FIG. 1:**
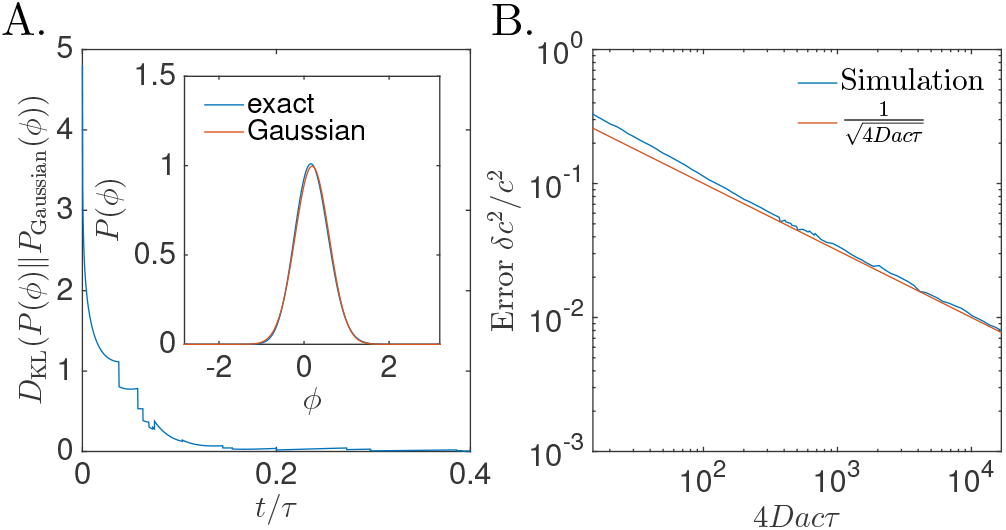
Numerical validations of analytical results. **A**. The Gaussian Ansatz (7)-(8) is validated by simulating the general equations for Bayesian filtering (5)-(6). The numerical solution approaches the Gaussian solution rapidly, as indicated by the decay of the Kullback-Leibler divergence 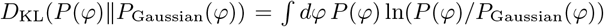. We used *rτ* = *Dacτ* = 50. **B**. Concentration sensing error as a function of concentration. The error estimated from simulations follows closely the prediction from (13), which is expected to be valid for 4*Dacτ* ≫ 1.

## Error estimate

To study the typical behaviour of (7)-(8), we now assume that the rate of binding events is large compared to the rate of change of the concentration, 4*Dacτ* = *rτ* ≫ 1. This regime is the biologically relevant one: to sense concentration, cells need to record many binding events over the time scale on which the concentration fluctuates. In that limit the estimator 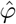 is close to the true value *φ*\*, and the Bayesian uncertainty *σ*^2^ is small, allowing for two simplifications. First, (8) relaxes over time scale *r*(*t*)^−1^ to a quasi-steady state value 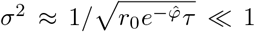. Second, we can make a small noise approximation for binding events: over some time interval Δ*t*, with *r*\*(*t*)^−1^ ≪ Δ*t* ≪ *τ*, the number of binding events has both mean and variance equal to *r*\*(*t*)Δ*t*, allowing us to replace discrete jumps in (7) by:

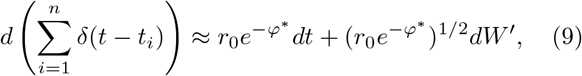

where *W*′ is a Wiener process. As a result, the estimator 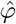 tracks the true value *φ*\* according to:

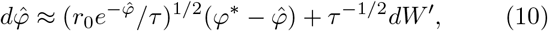

where we have expanded at first order in 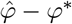. In the general case, the true field may evolve according to a different characteristic time scale, *τ*\*, than the one assumed by the Bayesian filter, *τ*, so that *dφ*\* = (*τ*\*)^−1/2^*dW*. The estimation error 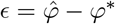 then evolves according to:

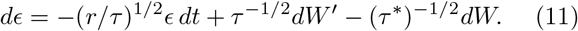

Intrigingy, the noises *dW*′ and *dW* have very different interpretations, one being due to the random arrival of binding events, and the other to the geometric diffusion of the concentration. Yet they come in the same form in this equation. Relying again on the assumption that *rτ* ≫ 1, we get an estimate of the error:

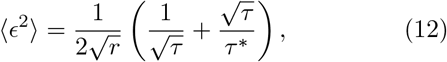

which has a minimum as a function of *τ*, reached for the true value of the characteristic fluctuation time *τ* = *τ*\*:

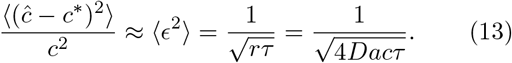

This error is equal to the Bayesian uncertainty 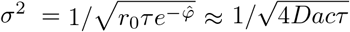 and is consistent with the error found using the saddle-point approximation in the related problem of probability density estimate [15].

We checked the validity of our small-noise approximation by comparing the prediction from (12) with the results of a numerical simulation of (7)-(8), in which we averaged the error 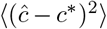 as a function of *c* for many realization of the process. The agreement is found to be excellent, and gets better as *rτ* = 4*Dacτ* becomes larger (Fig. 1B).

The error in (13) sets a fundamental physical limit on any concentration sensing device, biological or artificial, in a concentration profile that follows a geometric random walk. This bound is radically different from that obtained by Berg and Purcell for the concentration sensing error by a single receptor integrating binding events over time *T* [1, 5]:

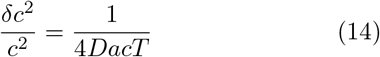

(in the limit where binding events are short so that the receptor is always free).

The major difference is that Berg and Purcell, as well as most of the literature on concentration sensing, assume that the sensed concentration does not change with time. Our result can be reconciled with Berg and Purcell by defining an effective measurement time 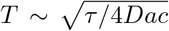 — the geometric mean between the mean time between binding events and the time scale of variation. This *T* realizes the optimal tradeoff between the requirement to integrate over many binding events, *T* ≫ 1/(4*Dac*), but over a relatively constant concentration, *T* ≪ *τ* [21].

## Plausible biological implementation

Can cells implement the optimal Bayesian filtering scheme and reach the bound set by (13)? To gain intuition, it is useful to rewrite (7)-(8) in term of the concentration estimator 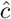, in the limit 4*Dacτ* ≫ 1 where *σ*^2^ can be eliminated:

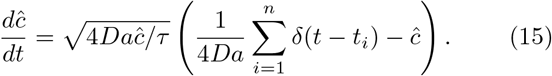

Each binding event should lead to an increment of 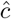, followed by a continuous, exponential decay, with a rate given by 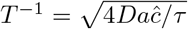.

This scheme can be implemented by a simple biochemical network schematized in Fig. 2A. The concentration readout 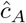 may be represented by the “active” (for instance phosphorylated) form *A*\* of a chemical species. Binding events cause the receptor to activate *A* into *A*\*, which gets subsequently deactivated. Both the activation and deactivation of *A* are catylized by a second chemical species in its active form, *B*\*. Thus, upon a binding event, the concentration of active *A*\* is increased by:

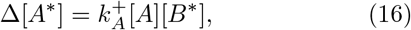

and it decays between binding events according to:

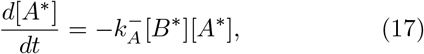

where 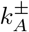 are biochemical parameters.

**FIG. 2:**
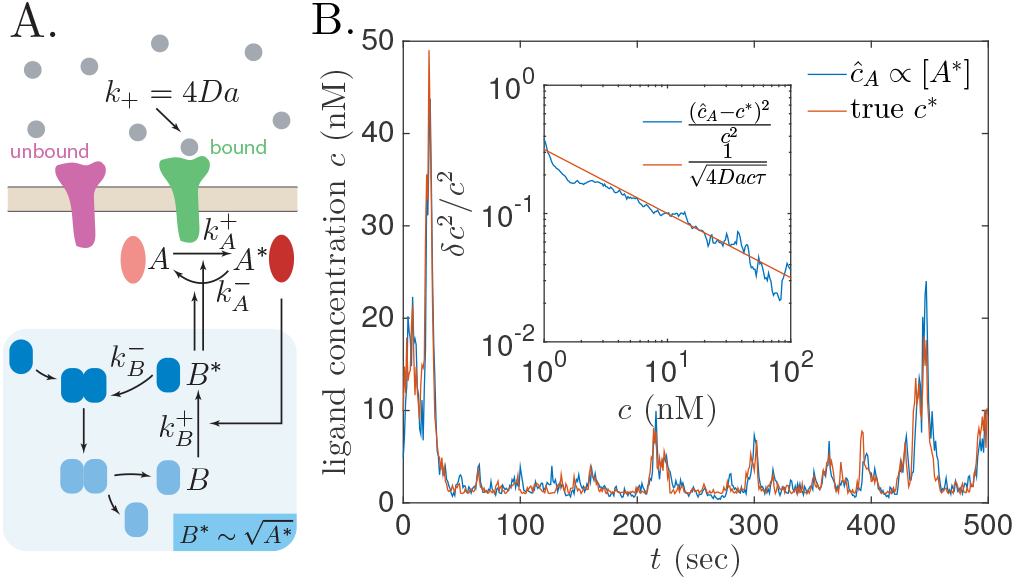
Performance of adaptive biochemical network in fluctuating ligand concentration. A. Schematic of the biochemical network implementing optimal Bayesian filtering. The receptor-induced activation of the readout molecule *A*\*, as well as its deactivation are regulated by a second molecule *B*\*, which is made to scale like 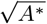 using a mechanism of deactivation by dimerisation (shaded box). **B**. Simulation of the network readout *c_A_*(*t*) ∝ *A*\* (*t*) in response to stochastic binding events in a fluctuating concentration field *c*\*(*t*). The relative estimation error 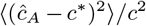 behaves according to the theoretical bound 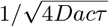 (inset).

To implement (15), the concentration of *B*\* must be controled by the square root of *A*\*. This dependence can be achieved by assuming that *B* is activated into *B*\* through the catalityc activity of *A*\*, and that *B*\* gets deactivated cooperatively as a dimer:

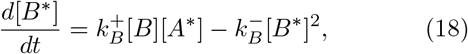

where 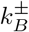 are biochemical reaction rates.

Assuming that the kinetics of *B* are fast compared to A, we obtain 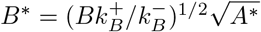 and

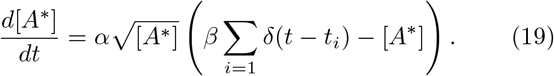

with 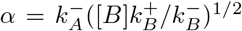 and 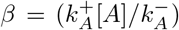. If *A* and *B* are in excess, and thus approximately constant, then this biochemical network exactly implements (15), with 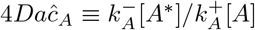, and 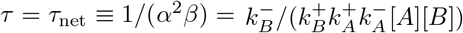.

Interestingly, the amount of inactive (≈ total) *B* controls the time scale of concentration fluctuations, and could be tuned through gene regulation to adapt to different speeds of environmental fluctuations. A biochemical network might be able to find the optimal *τ* and then adjust [B] accordingly by empirically measuring the fold-change of *r*(*t*) (which can be done by biochemical networks, see e.g. [22]) but with a delay, 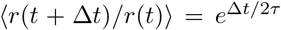, and then inverting the relationship to extract *τ*.

We tested the performance of the biochemical network for sensing concentration by simulating (16)-(18) with a fluctuating ligand concentration *c*(*t*) with characteristic time scale *τ*. For concreteness, we set *c*\*(0) = 10nM, *τ*\* = 10s, 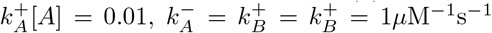 and [*B*] = 10*μ*M, so that *τ*_net_ = *τ*\*. Fig. 2B shows the network estimate 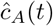 along with the true value *c*\*(*t*). The empirical error 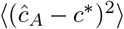 as a function of *c*\* averaged over 10^4^s (Fig. 2B, inset), again shows an excellent agreement with the theoretical bound 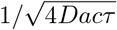.

## Discussion

For the sake of clarity our analysis made simplifying assumptions which can be easily relaxed. Our proposed biochemical implementation assumed a constant burst of activity following each binding event, consistent with the optimal estimation strategy. However, in real receptors, stochasticity in the bound time is known to double the variance in the estimate [5] (App. D [20]). Treating this effect simply adds a factor 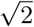 in the noise term of (9) as well as in (13), 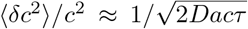. We also ignored periods during which the receptor was bound. During that time the receptor is blind to the external world, and the posterior evolves according to the prior: 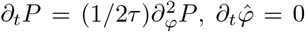 and 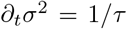. In our results, these “down times” renormalize the effective observation time by the fraction of time the receptor is free, *p*_free_ = (1+4*Dacu*)^−1^, where *u* is the average bound time, 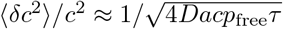 (App. D [20]). Combining the two effects (stochasticity in bound time and receptor availability) would yield 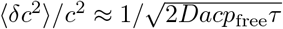.

The field theory of (2) is mathematically similar to the problem of estimating a density function from a small sample set with a smoothing prior [15–17, 19]. The main difference lies in the domain of observations. In density estimation the whole function {*φ*(*t*)}_*t*∈[0,*T*]_ is infered together on the whole domain of *t*, while sensors can only learn from past observations, *i.e*. the *t*′ < *t* half-plane. However, our solution can easily be generalized to deal with the entire time domain using the forward-backward algorithm (App. E [20]). Eqs. (5)–(6) and (7)–(8) can be solved both forward (from 0 to *t*) and backward (from *T* to *t*, with time reversal) in time, giving 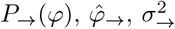 for the forward solution (the one treated in this article), and 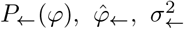 for the backward solution. The Bayesian posterior at any given time is then given by 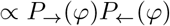, of mean 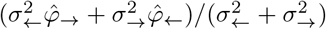 and variance 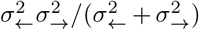 in the Gaussian approximation. While this situation is not relevant for concentration sensing, our general solution should be applicable to problems of density estimation. The saddle-point approximation usually made in that context [15–17] is expected to work in the same limit as our Gaussian Ansatz; however, recent work has emphasized the importance of non-Gaussian fluctuations for small datasets [19].

The biological implementation we propose is speculative. An interesting direction would be to identify square-root or similar control of receptor signaling in real biological systems, and interpret them in terms of optimal Bayesian filtering. Signaling pathways dealing with concentration changes over several orders of magnitude, such as bacterial chemotaxis, typically use adaptation mechanisms to increase the dynamic range of sensing [23]—a feature that is absent from our approach as we neglect noise in the signaling output. Combining adaptation design with ideas from Bayesian estimation could help us gain insight into the fundamental bounds and resource allocation tradeoffs that limit biological information processing.

## Acknowledgments

The authors would like to thank the Casa Matemática Oaxaca from the Banff International Research Station where this work was initiated. TM was partially supported by Agence National pour la Recherche (ANR) grant No. ANR-17-ERC2-0025-01 “IRREVERSIBLE” and IN by NSF Grants No. PHY-1410978 and IOS-1822677.

## Appendix A: Field theory for concentration sensing

We first recall the problem outlined in the main text for self-consistency. Receptor binding happens with rate *r*(*t*) = 4*Dac*(*t*) = *r*_0_*e*^−*φ*(*t*)^. The field *φ*(*t*) follows a random walk with characteristic time *τ*\*:

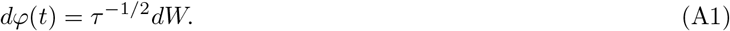

In the following *φ*\* will refer to the actual realization of the random walk, with characteristic time *τ*\*, and *φ* will refer to the guess made based on the observation of binding events, while *τ* denotes the assumes characteristic time scale.

The probability of the time trace of *φ* in the absence of any observation is given by

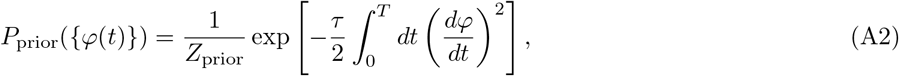

with *Z*_prior_ = (2*πdt*/*τ*)^*T*/2*dt*^, where *dt* is an infinitesimal discretization scale.

During each interval [*t*,*t* + *dt*] without a binding event, the likelihood reads *e*^−*dtr*(*t*)^ = *e*^−*dtr*_0_*e*^−*φ*(*t*)^^. A binding event in interval [*t*,*t* + *dt*] has likelihood *dtr*_0_*e*^−*φ*(*t*)^. Thus the posterior probability thus reads:

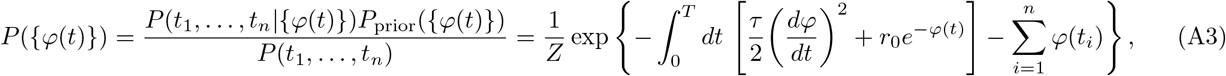

We define:

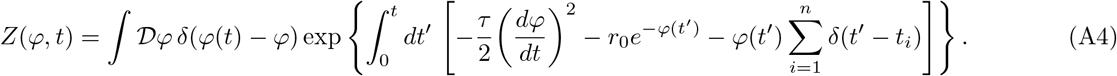

The marginal of *φ* at time *t*, *P*(*φ*, *t*|{*t*_1_, …, *t*_*n*′_}), where *n*′ is the last binding event before *t*, is then:

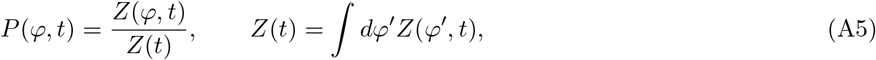

and *Z* = *Z*(*T*).

The partial partition function *Z*(*φ*, *t*) can be computed recursively. Let us start with the case where no binding occurs between *t* and *t* + *dt*. Then:

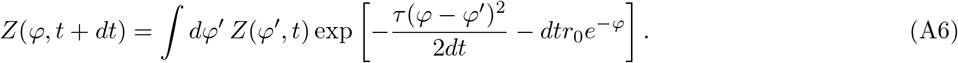

Equivalently,

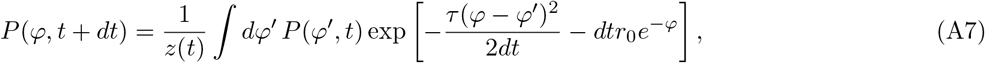

where

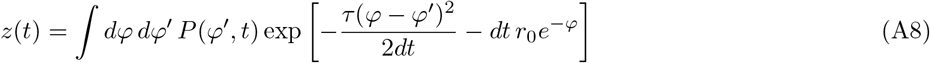

is a normalization constant, which can be used to calculate *Z*(*t*) recursively: *Z*(*t* + *dt*) = *Z*(*t*)*z*(*t*). Let us now take the limit *dt* → 0. The Gaussian integral becomes infinitely peaked. Expanding *P*(*φ*′, *t*) = *P*(*φ*, *t*) + *∂P*(*φ*, *t*)/*∂φ*(*φ*′ – *φ*) + *∂*^2^*P*(*φ*, *t*)/*∂φ*^2^(*φ*′ – *φ*)^2^/2, we obtain:

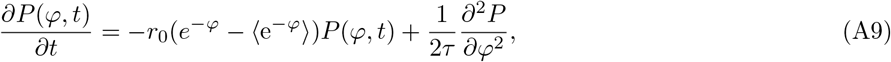

where 〈*f* (*φ*)〉 = ∫ *dφ P*(*φ*, *t*)*f*(*φ*), and

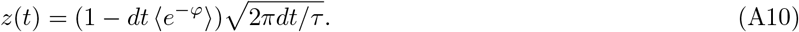

Defining 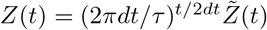, we get:

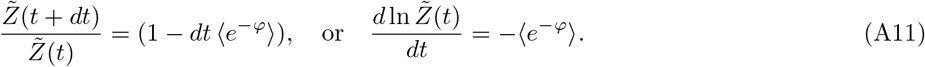

Now assume that there is a binding event between *t* = *t_i_* and *t_i_* + *dt*. Then *P*(*φ*, *t*) is discontinuous at *t_i_* and terms of order *dt* can be ignored:

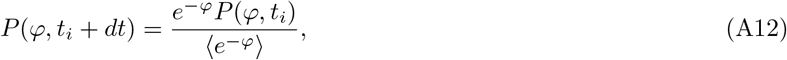

and 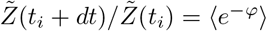.

In summary, the evolution equation for *P* reads:

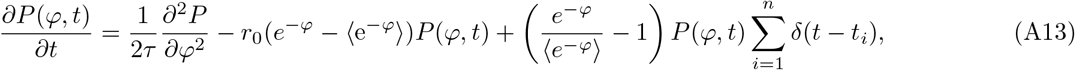

and the partition function is given by

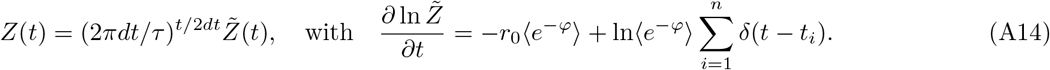

## Appendix B: Gaussian approximation

Because of the term 〈*e*^−*φ*^〉, (A13) is non-linear and cannot be solved analytically. However, if we assume that *P*(*φ*, *t*) is Gaussian,

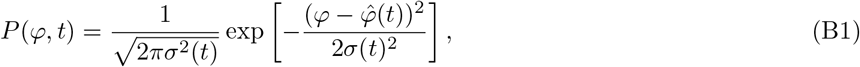

then closed equations can be obtained for the mean 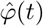 and variance *σ*^2^(*t*):

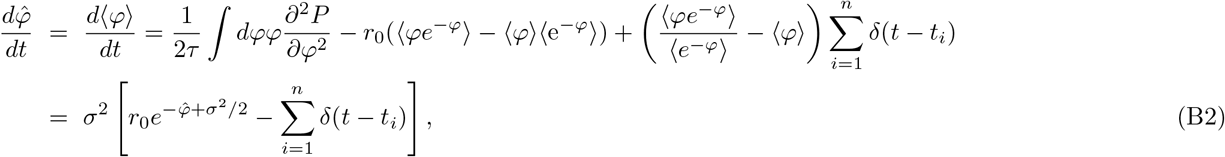

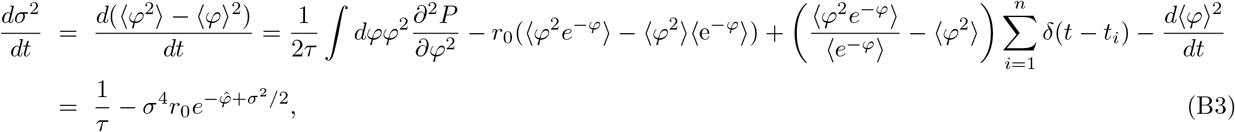

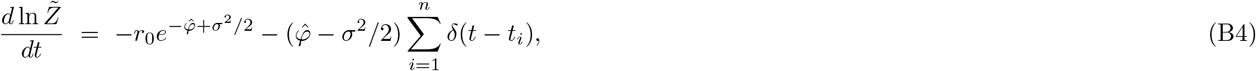

where we have used the Gaussian integral rules: 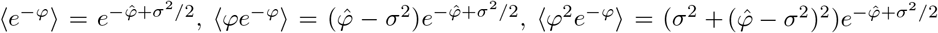, and 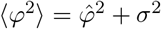, and we have used integration by parts to calculate the integrals.

## Appendix C: Partition function and time scale inference

The most likely timescale *τ* can be inferred from the observations as well by using Bayes’s rule again:

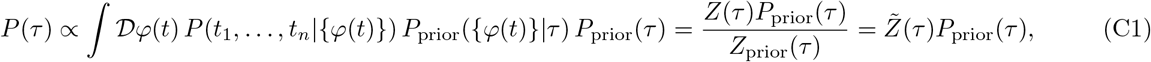

where we have used *Z*_prior_ (*τ*) = (2*πdt*/*τ*)^*T*/2*dt*^.

We can calculate 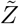 from the Gaussian approximation (B4):

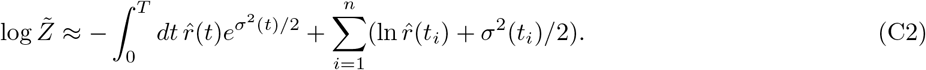

This expression looks like the log-likelihood of a sequence of binding events, up to the *σ*^2^ corretions. Bear in mind that 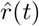 is the estimated rate, not the true one *r*\*(*t*). We have 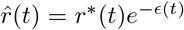, where we 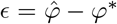. Expanding in e and *σ*^2^, we obtain:

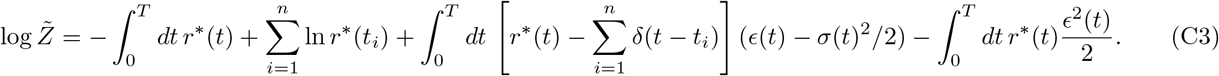

Both *ϵ*(*t*) and ∑_*i*_ *δ*(*t* – *t_i_*) – *r*\* are stochastic processes of mean 0. They are also uncorrelated with each other, so that the third term is sub-linear in *T*. The last term, which scales with *T*, thus dominates the *τ*-dependent part of the likelihood. It is maximized for minimum mean squared error, that is for *τ* = *τ*\*, as shown in the main text. At large *T*, the 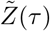 term exponentially dominates the prior *P*_prior_(*τ*), so that *P*(*τ*) is peaked around the maximum of 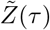.

Eq. (C2) is an example of the usual bias-variance tradeoff (where “bias” refers to errors made from overfitting the data, and “variance” to errors due to limited data). At small *τ*, 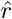 changes rapidly, jumping when a new binding happens, and then rapidly decreases. Thus the term 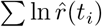 increases, indicating increase in the goodness of fit. At the same time *σ*^2^(*t*) increases, so that at small *τ* the integral term in (C2) becomes large and negative, exploding exponentially for *τ* → 0. In contrast, for *τ* → ∞, 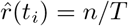, and *σ*^2^ → 0, so that now the goodness of fit is small.

Overall, theres an optimal *τ* that maximizes 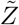. The three terms in (C2) parallel the three terms (fluctuation determinant, goodness of fit, and the kinetic term) in the field-theoretic formulation of continuous probability density estimation from samples [15, 16], and thus we expect that maximizing 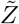 will result in an optimal *τ* not only when the true concentration undergoes a geometric random walk, but also when it undergoes various anomalous walks [16].

## Appendix D: Bound time

We have so far neglected the time the receptor remained bound the ligand Let us denote by *t*_*i*, off_ the unbinding time following the binding time at *t_i_*. When the receptor is bound, *t_i_* < *t* ≤ *t*_*i*, off_, no information can be obtained from the environment, and the evolution equation for the posterior simply follows the diffusion law:

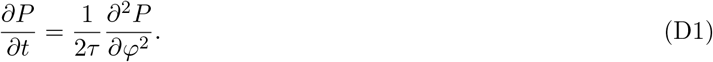

The rest of the time, *t*_*i*, off_ < *t* ≤ *t_i_*, (A13) holds. Similarly, in the Gaussian approximation, we get

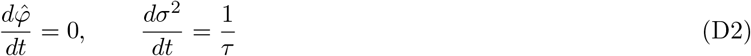

for bound receptors, *t_i_* < *t* ≤ *t*_*i*, off_, and (B2)-(B3) for unbound receptors. In the limit where binding and unbinding events are frequent compared to *τ*, *rτ* ≫ 1, we have:

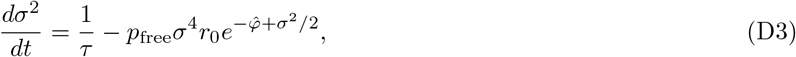

where

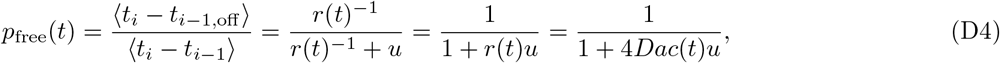

where *u* = 〈*t*_*i*, off_ – *t_i_*〉 is the average bound time, and *r*(*t*) ^1^ = 〈*t_i_* – *t*_*i*–1, off_〉 the average unbound time. The uncertainty then reads:

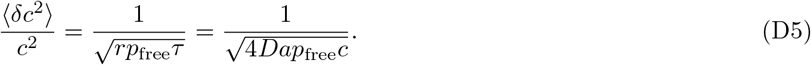

The time the receptor remains bound has another impact on the ability to sense concentration. In the biochemical scheme proposed in the main text, each binding event causes a fixed burst of activity *δ*(*t* – *t_i_*), regardless of the bound time. In the simplest receptors however, signaling occurs during the time the receptor is bound, which is itself stochastic. We can model this by replacing the Dirac delta by a random burst of activity, *b_i_δ*(*t* – *t_i_*), with *b_i_* proportional to the bound time, *b_i_* = (*t*_*i*, off_ – *t_i_*)/*u*, so that 〈*b_i_*〉 = 1 and Var(*b_i_*) = 1, since the bound time is distributed exponentially according to (1/*u*)*e*^−(*t*_*i*, off_–*t_i_*)/*u*^. More generally we can consider 〈*b_i_*〉 = 1 and Var(*b_i_*) = *CV*, where 0 ≤ *CV* ≤ denotes the coefficient of variation. The general base 〈*b_i_*〉 can be renormalized away into the biochemical parameters. The special case *CV* = 0 gives back the results of the main text. When *CV* > 0, the variance of 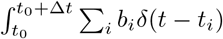 over an interval of duraction Δ*t* instead reads:

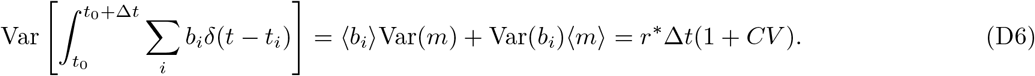

where *n*′ is the number of binding events in the interval. As the result, the noise *dW*′ in the main text gains a factor 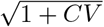, and the error becomes:

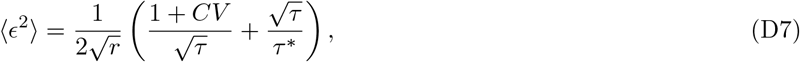

and minimal error reached for *τ* = (1 + *CV*)*τ*\*:

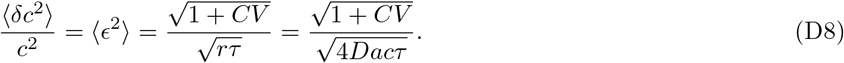

Taking into account both receptor occupancy and stochasticity in bound times finally yields:

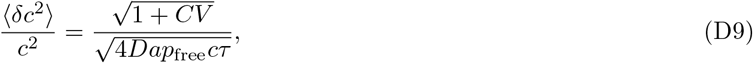

or, in the case of complete stochastic unbinding, *CV* = 1:

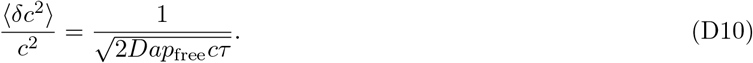

## Appendix E: Beyond concentration sensing – using the future

Information about future binding events can be exploited by using the backward equation for *P*_←_ = *P*(*φ, t*|{*t*_*n*′+1_, …, *t_n_*}):

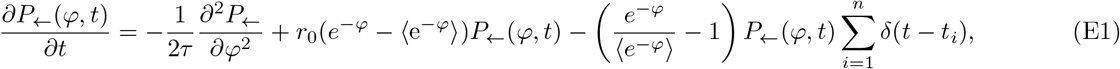

where *m* is the last binding event before *t*. We denote by *P*_→_ = *P*(*φ, t*|{*t*_1_, …, *t*_*n*′_}) the solution of the forward equation (A13) discussed before. The distribution of *φ* at time *t* is then given by:

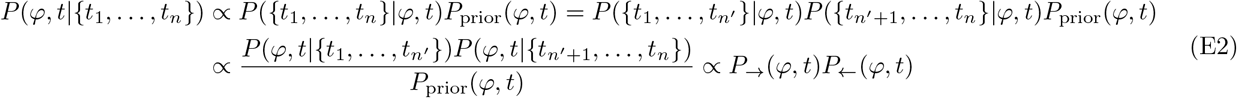

where we have used the fact that the past and future where conditionally independent given *φ* at time *t*, since the process is Markovian, and a uniform prior. We thus have:

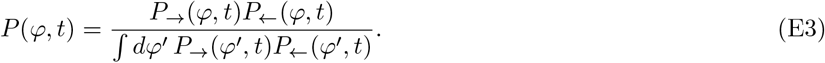

Using the Gaussian solution, and denoting 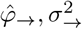 the parameters of the forward solution, and 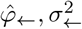 those of the backward solution, we obtain:

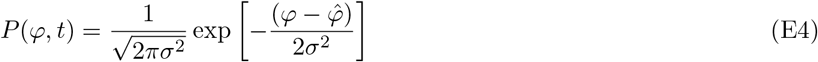

with

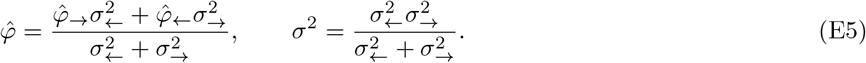

These formulas could be used in estimates of density *r*(*t*) from sparse observations *t_i_*, where *r*(*t*) is interpreted as a density of events, and *t* is the variable whose density we want to infer.

## References

[1] H. C. Berg and E. M. Purcell, Biophys. J. 20, 193 (1977).

[2] W. Bialek and S. Setayeshgar, Proc. Natl. Acad. Sci. U. S. A. 102, 10040 (2005).

[3] K. Kaizu, W. De Ronde, J. Paijmans, K. Takahashi, F. Tostevin, P. R. T. Wolde, and P. R. ten Wolde, Biophys. J. 106, 976 (2014).

[4] G. Aquino, N. S. Wingreen, and R. G. Endres, J. Stat. Phys. 162, 1353 (2016).

[5] R. G. Endres and N. S. Wingreen, Phys. Rev. Lett. 103, 158101 (2009).

[6] E. D. Siggia and M. Vergassola, Proc. Natl. Acad. Sci. U. S. A. 110, E3704 (2013).

[7] J.-B. Lalanne and P. Francois, Proc. Natl. Acad. Sci. 112, 1898 (2015).

[8] T. Mora, Phys. Rev. Lett. 115, 038102 (2015).

[9] V. Singh and I. Nemenman, PLoS Comput. Biol. 13, 1 (2017).

[10] V. Singh and I. Nemenman, arXiv:1906.08881 (2019).

[11] M. Carballo-Pacheco, J. Desponds, T. Gavrilchenko, A. Mayer, R. Prizak, G. Reddy, I. Nemenman, and T. Mora, Phys. Rev. E 99 (2019).

[12] E. Kussell and S. Leibler, Science 309, 2075 (2005).

[13] T. Mora and N. S. Wingreen, Phys. Rev. Lett. 104, 1 (2010).

[14] Z. H. E. Chen, Statistics 182, 1 (2003).

[15] W. Bialek, C. Callan, and S. Strong, Phys. Rev. Lett. 77, 4693 (1996).

[16] I. Nemenman and W. Bialek, Phys. Rev. E 65, 2 (2002).

[17] J. B. Kinney, Phys. Rev. E - Stat. Nonlinear, Soft Matter Phys. 90 (2014).

[18] J. B. Kinney, Phys. Rev. E - Stat. Nonlinear, Soft Matter Phys. 92 (2015).

[19] W. C. Chen, A. Tareen, and J. B. Kinney, Phys. Rev. Lett. 121, 160605 (2018).

[20] See Supplemental Material for detailed derivations.

[21] A. Kolmogoroff, C. R. Acad. Sci. Paris 208, 2043 (1939).

[22] L. Goentoro, O. Shoval, M. W. Kirschner, and U. Alon, Mol. Cell 36, 894 (2009).

[23] M. D. Lazova, T. Ahmed, D. Bellomo, R. Stocker, and T. S. Shimizu, Proc. Natl. Acad. Sci. U. S. A. 108, 13870 (2011).

